# Incidence of atrial septal defects in children attended the Cardiac Center of Ethiopia during January 2016 to December 2018

**DOI:** 10.1101/2020.02.12.945360

**Authors:** Abdu Hassen Musa, Mekbeb Afework, Mohammed Bedru, Shibikom Tamirat, on behalf of Cardiac Center of Ethiopia record screener

## Abstract

**Background:** Although it appears that an atrial septal defect (ASD) occurs frequently in Ethiopia there are only a few published studies available so far on this matter. This study is therefore aimed to evaluate the prevalence and echocardiographic characteristics of ASDs in children (aged ≤ 15 years) attended a cardiac referral center in Ethiopia.

**Methods:** This retrospective study reviewed the sociodemographic data and the initial echocardiographic findings of the children with ASDs who were diagnosed at the Cardiac Center of Ethiopia (CCE), Addis Ababa, from January 2016 to December 2018.

**Results:** A total of 116 children (56.9% females and 43.1% males) with a mean age of 3.47± 3.72 years (range: 15 days to 15 years) were diagnosed with ASDs. The most prevalent age groups were infancy (50%) and early childhood (29.3%). All the studied cases were diagnosed with ostium secundum ASD while there were no cases with ostium primum, sinus venosus and coronary sinus defects. The most frequent ostium secundum ASD was large size (61.2%) and was more frequent in infants (23.3%) and young children (21.6% of all cases).

**Conclusion:** Ostium secundum ASD is the most prevalent IAS defect and more common among female cases. Large size ostium secundum ASD is more frequent in the studied children and is more prevalent in infants and young children. This survey may provide data for the currently lacking statistics on ASDs at the CCE and might be helpful for the management and follow-up of children with ASDs. Scheduled follow-up and intervention studies are required to evaluate the incidence and patterns of spontaneous and surgical closure of ASD and their outcomes.

## Introduction

An ASD is an opening (a hole) in the interatrial septum (IAS) that separates the right and left atria. This opening results in interatrial communication or causes the mixing of blood from the systemic and pulmonary venous circulations. The ASD may occur in isolation or in association with other cardiac diseases such as tricuspid regurgitation, pulmonary stenosis, patent ductus arteriosus, and pulmonary arterial hypertension [1-3]. The ASDs can be classified as morphologic defects of the IAS and as defects that do not involve the IAS. Morphologic defects of the IAS could be ostium primum and/or ostium secundum defects. On the other hand, defects that do not involve the IAS could behave physiologically like interatrial septal defects or abnormal openings in the cardiac terminations of the systemic and pulmonary veins. This could indirectly cause communications of the right and left atria, and may include sinus venosus and coronary sinus defects [4-6].

The ostium secundum defect is the most common form of ASD. It represents about 75– 93.2% of ASD and can be seen as a defect of variable size in the region of the fossa ovalis. It is bounded by the superior, anterior, posterior, and inferior limbus of the fossa ovalis [6-8]. Majority of the ostium secundum defects are identified as single defect, while in some cases, are seen as multiple (fenestrated) defects [1,5,7].The ostium primum defect is considered as the mildest form of ASD, accounting for 15 – 20% of all ASDs. The primum ASD is located in the lower portion of the septum primum or in the atrioventricular (AV) canal portion of the interatrial septum. In the ostium primum defect, the AV valves are almost always abnormal and incompetent [4, 9,10].

The sinus venosus defects represent approximately 5 to 10% of all ASDs. This defect is unrelated to the interatrial septum. A sinus venosus defect is characterized by lack of septation between the right pulmonary veins and the superior vena cava (SVC) or right atrium. There are superior and inferior types of sinus venosus ASDs [8, 10,11]. The superior sinus venosus ASD (the most common form, accounting for 87% of sinus venosus defects) is a communication between the right upper pulmonary vein and the cardiac end of the SVC. The inferior sinus venosus defect is a communication between the right lower pulmonary vein and the right atrium just above the inferior vena cava-right atrial junction [4,8,10]. The coronary sinus defect is one of the rare forms of atrial communication resulting from partial or complete un-roofing of the tissue separating the coronary sinus from the left atrium. In the absence of other major structural anomalies, blood from the left atrium enters the coronary sinus and drains into the right atrium through the coronary sinus defect, which is typically enlarged to accommodate the increased blood flow [4,12].

Several studies have documented that ASD can close spontaneously (naturally) without surgical intervention, depending on the anatomical type and size of the defects, as well as age of the patients [13-15]. Size of the defect is the most important predictor for spontaneous closure of ASD [14,16]. On the other hand, sinus venosus and ostium primum defects are usually associated with a haemodynamically significant shunt, do not decrease in size, and usually need surgical closure [17, 18]. However, spontaneous closure of secundum defects has been frequently reported in young patients [17, 18]. A small size secundum ASD has shown spontaneous closure in 66 to 80% of cases, who were diagnosed within the first two years of life [14,15,18].

Closure of moderate secundum ASDs have been reported to occur spontaneously in 4 - 14% of young patients [13,14, 16]. However, several studies have consistently documented lack of spontaneous closure of large size ASDs which most often require surgery [6,16,19]. In spite of the spontaneous closure of ASDs, some studies have also noted an increased and/or a decreased size of ASDs, over time. For example, in follow up studies of children with moderate to large size ASDs and remained untreated, 65 to 75% showed increased, 9 to 14% showed reduced, while 15 to 20% showed no size changes in the diameter of ASDs, as their age increased [4,14,16,19].

The size of the ASD determines the pathophysiology of the defect. A small ASD often produces no apparent symptoms, because most small ASDs are associated with a fairly small shunt and with a minimum magnitude of blood flow through the defect. Thus, most children with small size defects are asymptomatic throughout their childhood, and sometimes the defects may be undetected until adulthood, due to lack of prominent clinical symptoms [20-22].

Moderate or large ASDs are associated with significant left-to-right shunt which could trigger a cascade of changes in the myocardium and in the pulmonary vasculature [23,24] and produce sign and symptoms[2,25]. Children with moderate or large ASDs may present with tachypnoea, slow weight gain, or recurrent respiratory infections. Over time they may complain shortness of breath, exercise intolerance (dyspnea and fatigue), palpitations, syncope, peripheral oedema and cyanosis [22,26,27].

Most often, ASD is sporadic or with no identifiable causes [1]. However, chromosomal abnormalities (e.g. mutations in the cardiac transcription factors that are important during atrial septation including gene *NKX2-5, GATA4* and *TBX5* located on chromosome 14q12) and maternal exposure to certain teratogenes, such as maternal alcohol and cigarette consumption, during pregnancy have been linked to ASDs [28-33].

ASD represents the second most common congenital heart diseases in pediatric populations, next to the ventricular septal defect. It was estimated to account for 7 to 23.7% of all CHDs, as reported in the previous studies done in Egypt [34], Saud Arabia [35], Ethiopia [36], and in USA [37]. The overall prevalence of ASD was estimated at 2 to 3.89 per 1000 children [4,5,38-40]. ASD is more common among female cases. An ostium secundum is the most common type of ASD; accounting for 75– 93.2% of all ASD and females constitute 65–75% of patients with the secundum ASD [4,5,39,40]. At the time of this write-up, there are no published studies and statistics available regarding the ASDs, in Ethiopian children. Majority of the available reports were from studies of congenital heart diseases [36]. Therefore, this survey was designed to document the incidence and echocardiographic profiles of ASDs in children who attended in the cardiac referral center of Ethiopia (Cardiac Center of Ethiopia). This study may provide a baseline data for the currently lacking statistics and for future studies on ASDs. Moreover, it may help in the management and follow-up of children with ASD.

## Material and methods

This was a retrospective study and retrieved echocardiographic reports, which were performed at the Cardiac Center of Ethiopia (CCE) during January 2016 to December 2018. The CCE is a referral center for cardiovascular care in Ethiopia, which is located at Addis Ababa city inside Black Lion Specialized Hospital. The center has pediatric and adult cardiologists, cardiac surgeons, and anesthesiologists. It also has facilities for electrocardiography echocardiography. The CCE receives cardiac patients referred from governmental and other health institutions of the country.

The study population consisted of children between the ages of 0 to 15 years who attended the CCE for various cardiac diseases, as of January 2016 to December 2018. During the period of review, a total of 7,452 patients were diagnosed with echocardiography, of these, 6341 patients aged > 15 years were excluded from ASD assessment. Hence, the remaining 1,111 pediatric patients (≤15 years) were included in the primary data abstraction. Of these children, a total of 995 cases with complex congenital heart diseases, valve heart diseases, ventricular septal defects and those with non ASD cardiac structural abnormalities were excluded from further data analysis, while, the remaining 116 children (≤ 15 years), that were diagnosed with ASDs were included and their clinical data were analyzed for the present study.

Data including age, sex, geographic origin and echocardiographic findings were collected, by record screener (nurses at the CCE) and medical reviewer (principal investigator), using a structured checklist. To avoid repetition bias, only the first time echocardiographic report of each pediatric case was utilized for data collection.

The classification and measurement of the ASD was based on the guidelines for the echocardiographic assessment of atrial septal defect and patent foramen ovale of the American Society of Echocardiography and Society for Cardiac Angiography and Interventions [4]. Data were entered and processed with a computer using SPSS version 20 software. Chi-square test was used to evaluate possible differences between proportions. The level of significance was set at p< 0.05. Permissions to carry out this study were obtained from the research and ethics committees of Anatomy Department and Cardiac Center of Ethiopia.

## Results

Over the three years study period, the initial echocardiographic reports of 1111 children were reviewed, of these, 116 (10.44%) children were screened with atrial septal defects. Out of the total 116children with ASD, 66 (56.9%) were females and 50 (43.1%) were males, with a female to male ratio of 1.32: 1.

At the time of presentation, the age range of the patients was 15 days to 15 years with a mean age of 3.47± 3.72 (M±SD) years. Of the total cases, 58 (50%) children were diagnosed within the first year of life, 34 (29.3%) were during toddler and early childhood (1 to 5 years), 15 (12.9%) were in middle childhood (5 to 10 years) and 9 (7.8%) cases were diagnosed during late childhood (> 10years), as depicted in Table 1.

**Table 1:** Frequency and percentage of patients with ASD by age and sex (N=116).

| Variables | Frequency and %age |  |  | P - value |
| --- | --- | --- | --- | --- |
|  | Female n (%) | Male n (%) | Total |  |
| Gender prevalence | 66(56.9%) | 50(43.1%) | 116 (100%) | 0.001* |
| ≤1 year | 29 (25%) | 29 (25%) | 58 (50%) | 0.001* |
| >1 to 5 years | 20 (17.2%) | 14 (12.1%) | 34 (29.3%) | 0.025* |
| >5 to 10 years | 9 (7.7%) | 6 (5.2%) | 15 (12.9%) | 0.241 |
| >10 years | 8 (6.9%) | 1(0.9%) | 9 (7.8%) | 0.341 |

*\*P value < 0.05 was considered statistically significant.*

As indicated in the Table 2, out of the total children with ASD, over the three years study period, significant (p< 0.05) numbers of children with ASD were from Addis Ababa city (52.6%), the surrounding Oromiya region (29.3%) and Amhara region (9.5%).

**Table 2:**
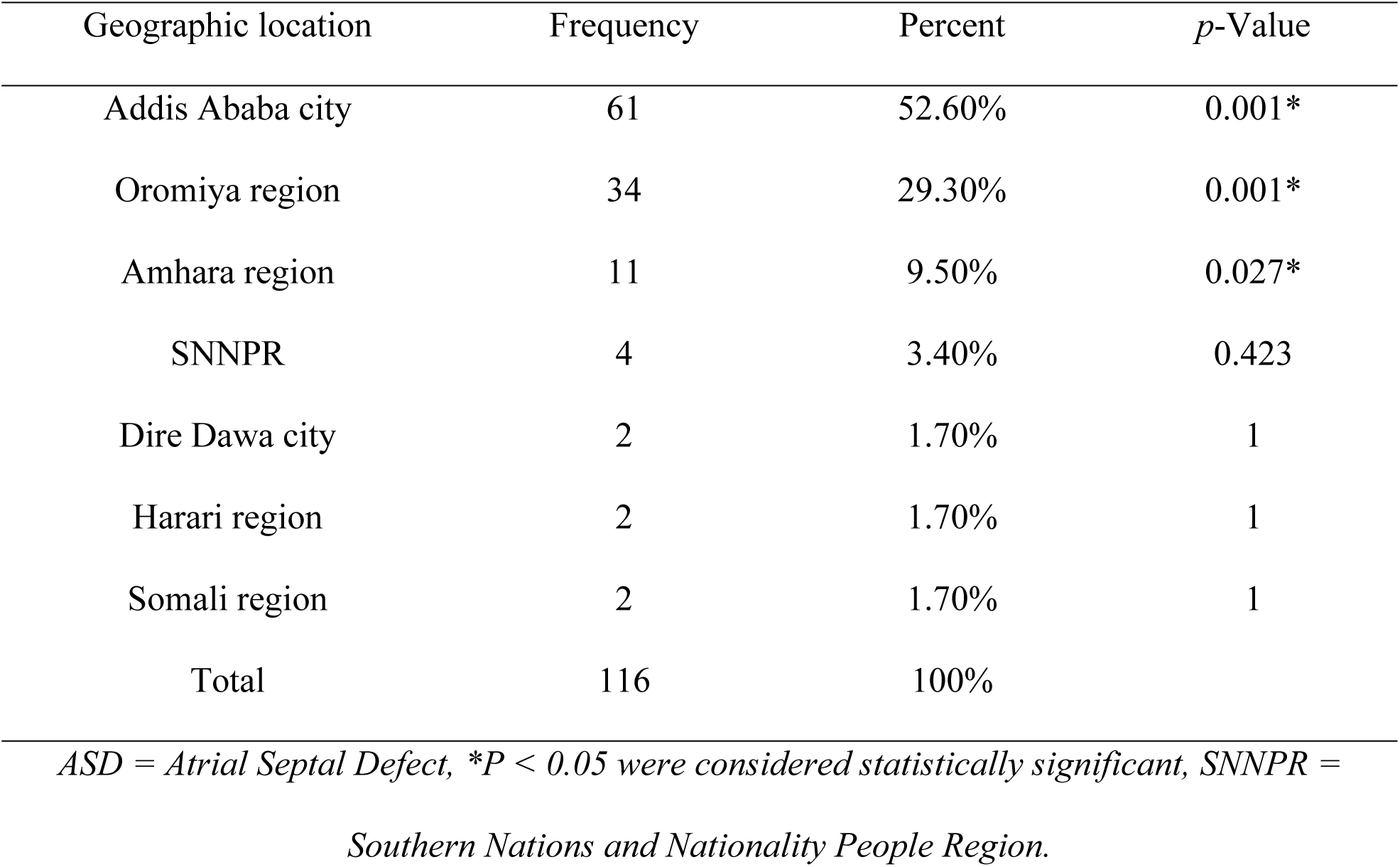
Geographic origin of the children with VSD (N=116).

Over the three years review period, all (100%) cases had ostium secundum ASDs (including, 96.6% with single and 3.4% with multiple secundum ASDs). There were no children diagnosed with ostium primum, sinus venosus and coronary sinus defects.

Of the total children with secundum ASD, 61.2% (71 cases) had large size, 30.1% (43 cases) had small size and 1.7% (2 cases) had moderate size defects. The large size secundum ASD was the most frequent defect and was seen in more than half (61.2%) of the total cases, followed by small size ASD (30.1%), while moderate size defect was less frequent among the studied cases. Moreover, large size defect was more common in patients who were diagnosed during infancy (23.3%) and in early childhood (21.6%), while small size defect was more frequent in infants (25.9%), as presented in Table 3.

**Table 3:** Distribution of secundum ASD by size and age group of patients (N=116).

| Size of ASD | Total cases | Age group of cases |  |  |  |
| --- | --- | --- | --- | --- | --- |
|  |  | ≤1year | >1-5years | >5-10years | >10-15years |
| Small | 43 (37.1%) | 30(25.9%) | 8(7%) | 2(1.7%) | 3(2.6%) |
| Moderate | 2 (1.7%) | 1(0.85%) | 1(0.85%) | 0(0) | 0(0) |
| Large | 71(61.2%) | 27(23.3%) | 25(21.6%) | 13(11.2%) | 6(5.2%) |
| Total | 116 (100%) | 58 (50%) | 34(29.3%) | 15(12.9%) | 9(7.8%) |

*ASD = Atrial Septal Defect,*

## Discussion

This retrospective study retrieved and analyzed the sociodemographic and echocardiographic data of the children with ASD who attended the Cardiac Center of Ethiopia from January 2016 to December 2018. The CCE is the only cardiac referral center in the country that receives cardiac patients across the country for further investigation and management.

Over the three years study period, a total of 1111 children were referred to the CCE and diagnosed with various cardiac diseases including ASD, ventricular septal defects, valvular heart diseases, patent foramen ovale, tetralogy of Fallot, patent ductus arteriosus, transposition of great arteries and other cardiac anomalies. Of the total cases who were diagnosed during the study period, 116 children were identified with ASDs. This indicated that the prevalence of ASD was 10.44%. This is in line with the prevalence of 7 to 15% reported in Atlanta [41], Taiwan [42], Germany [43], Worldwide [44], and in Iran [6].

More than half (52.6%) of the cases were from Addis Ababa city and around one third (29.3%) of the patients were from the surrounding Oromiya region, whereas a small number of cases were from Dire Dawa city (1.7%), Harari region (1.7%) and from Somali region (1.7%). Although further studies are needed, the reason for the higher proportion of patients from Addis Ababa city and from the surrounding Oromiya region might be due to accessibility of the cardiac center to these regions. However, it might also be associated with the life style of pregnant mothers living in larger cities (Addis Ababa and the surrounding Oromiya region) making them more exposed to teratogens (such as maternal alcohol consumption, cigarette smoking, drug exposure as well as maternal diabetes) than those from smaller cities and rural areas.

Several studies have suggested that the non-genetic factors such as poorly controlled maternal diabetes, maternal alcohol, drugs and cigarette consumptions during pregnancy are linked to the congenital heart disease including ASDs [28-33, 45].

Over the review period, out of the total children with ASD, 56.9% were females and 43.1% were males (with a female to male ratio of1.32: 1). Similar results where ASD was more common among female cases were reported in the previous studies done in Europe [46], India [47], Nigeria [48], and in Iran [6].

Out of the total children with ASDs, 50% were diagnosed at infancy, 29.3% were seen at early childhood, 12.9% were at middle childhood and 7.8% were at late childhood. It appears that, more than two third (79.3%) of the total cases with secundum ASD were diagnosed during infancy and early childhood (≤ 1 and > 1 to 5 years). This might be related to the findings of the higher incidence of the defect with severe or large size ASD in the studied cases and presentation of clinical symptoms at earlier ages. It was documented that, a small ASDs are commonly associated with a fairly small shunt and with a minimum magnitude of blood flow through the defect and often produces no apparent symptoms, because the heart and the lungs may not have to work harder than the usual [2,40,49,50]. Whereas large ASDs are haemodynamically significant and allow up to 2 to 3 times the ordinary amount of blood to circulate through the right chambers of the heart and the lungs. This makes the heart and the lungs to do more work than the usual. If the heart and the lungs are unable to cop-with the extra work, symptoms of congestive heart failure could develop.

Patients with moderate or large ASDs are characterized by marked signs and symptoms of congestive heart failure (including poor growth and development, respiratory distress, and recurrent respiratory infections). Over time, they may complain shortness of breath, exercise intolerance (dyspnea and fatigue), palpitations, syncope, peripheral oedema and cyanosis [2,22,49,51,52]. Presentation of these sign and symptoms due to large size secundum ASD may necessitate the early diagnosis in the current studied cases. The current finding is in line with the studies done in Europe [46], India [47], and in Iran [6].

During the study period all detected cases with ASD were diagnosed with ostium secundum defects (100%), with no children diagnosed with ostium primum, sinus venosus and coronary sinus defects. The current higher prevalence of secundum ASD is in agreement with the studies done in USA [7,8], Nigeria [48] and in Iran [6].

Of the total cases, the relative size of secundum ASDs was large in 61.2% (71 cases), small in 30.1% (43 cases) and moderate in 1.7% (2 cases). Of the 71 cases with large size defects, 38% (27 patients) were infants, 35.2% (25) were younger children, 18.3% (13) were in middle childhood and 8.5% (6) were older children. It seems that, large size secundum ASD was more frequent in infants and younger children representing 73.2% of all cases with large size ASD and 44.8% of the total cases.

The current higher incidence of large size secundum ASD and its preponderance in infants and younger children was in accordance with the previous studies conducted in Austria [14], Europe [46], India [47], and in Iran [6].

## Conclusion

The present study has revealed that secundum ASD was the commonest defect in studied cases and was more prevalent in females than males. It was also more frequent among infants and young children. Furthermore, large size secundum ASD was more prevalent in the studied cases and was more common during infancy and early childhood.

## Acknowledgment

We would like to acknowledge Dr. Helen Befikadu (medical director of the CCE) and Mr. Hiruyi Ali (manager of the CCE) for their support and permission. We wish to thank Mr. Biniyam Alemayehu and Miss Selamawit Nigusse (record screener at the CCE) for their active participation and support in the data collection. The authors also acknowledge Addis Ababa University and Dilla University for the financial support of the study.

## Reference

1. Kliegman, RM, Behrman RE, Jenson HA, Stanton BF. Nelson Textbook of Pediatrics, 18^th^ ed. Copyright © Saunders, An Imprint of Elsevier;2007.

2. Bonow RO, Mann DL, Zipes DP, Libby P. Braunwald’s heart disease: a textbook of cardiovascular medicine (^9th^ed.). Philadelphia Copyright © 2012 by Saunders, an imprint of Elsevier Inc.: International Edition. 2012; 978 –89.

3. Brunicardi FC, Anderson DK, Billiar TR, Dunn DL, Hunter JG, Matthews JB, et al. Schwartz’s Principles of Surgery; (^9th^ Ed). Copyright © The McGraw-Hill Companies; 2012.

4. Silvestry FE, Meryl SC, Laurie BA, Nitin JB, Craig EF, et al. ASE/SCA Guidelines for the Echocardiographic Assessment of Atrial Septal Defect and Patent Foramen Ovale: From the American Society of Echocardiography and Society for Cardiac Angiography and Interventions. Journal of American Society for Echocardiography. 2015; 28:910–58.

5. Adler DH, Ellis AR. Atrial septal defect. The heart org. Medescape-e Medicine. 2017; 1–10.

6. Behjati-ArdakaniM, Golshan M, Akhavan-Karbasi S, Hosseini SM, Sarebanhassanabadi M. The Clinical course of patients with atrial septal defects. Iran Journal of Pediatrics. 2016;26:4: 1–10.

7. Patel AR, DAlessandro L, Weinberg PM. Anatomy of the atrial septum. In: Hijazi ZM, Feldman T, Abdullah Al-Qbandi MH, Sievert H, editors. Transcatheter Closure of ASD and PFO. 1st ed. Minneapolis: Cardiotext Publishers. 2010; pp. 3–15.

8. Carlos A, Rojas CA, El-Sherief A, Medina HM, Chung JH, Choy G, et al. Embryology and Developmental defects of the interatrial septum. American Journal of Roentgenology. 2010; 195:1100–1104.

9. Kirby ML. Endocardium, cardiac cushions, and valve development. In: Cardiac Development. Oxford UK: Oxford University Press. 2007; pp. 119–31.

10. Banka P, Bacha E, Powell AJ, Geva T. Outcomes of inferior sinus venosus defect repair. Journal of Thoracic and Cardiovascular Surgery. 2011; 142:517–22.

11. Van Praagh S, Carrera ME, Sanders SP, Van Praagh R. Sinus venosus defects: unroofing of the right pulmonary veins—anatomic and echocardiographic findings and surgical treatment. American Heart Journal. 1994; 128: 365–79.

12. Ootaki Y, Yamaguchi M, Yoshimura N, Oka S, Yoshida M, Hasegawa T. Unroofed coronary sinus syndrome: diagnosis, classification, and surgical treatment. Journal of Thoracic Cardiovascular Surgery.2003; 126:1655–1656.

13. McMahon C, Feltes TF, Fraley J, et al. Natural history of growth of secundum atrial septal defects and implications for transcatheter closure. Heart Journal. 2002;87 (3):256–9.

14. Hanslik A, Pospisil U, Salzer-Muhar U, Greber-Platzer S, Male C. Predictors of spontaneous closure of isolated secundum atrial septal defect in children: a longitudinal study. Pediat. Jour. 2006; 118(4):1560–5.

15. Ozcelik N, Atalay S, Tutar E, Ekici F, Atasay B. The prevalence of interatrialseptal openings in newborns and predictive factors for spontaneous closure. International Journal of Cardiology. 2006;108 (2):207–11.

16. Demir T, Oztunc F, Eroğlu AG, et al. Outcome for patients with isolated atrial septal defects in the oval fossa diagnosed in infancy. J. Cardiol. 2008; 18(1):75–8.

17. Geva T, Martins JD, Wald RM, Atrial septal defects: Seminar. Lancet. 2014; 383: 1921–32.

18. Wang SY, Welch TD, Elfenbein A, Kaplan AV. Spontaneous Closure of a Secundum Atrial Septal Defect. Methodist Debakey. Cardiovascular Journal. 2018; 14 (1)18.

19. Azhari N, Shihata MS, Al-Fatani A. Spontaneous closure of atrial septal defects within the oval fossa. Cardiology of Young. 2004; 14(2):148–55.

20. Rigatelli G. Congenital heart diseases in aged patients: clinical features, diagnosis, and therapeutic indications based on the analysis of a twenty five-year Medline search. Cardiology Review. 2005; 13:293–296.

21. Saxena A, Divekar A, Soni NR. Natural history of secundum atrial septal defect revisited in the era of transcatheter closure. Indian Heart Journal. 2005; 57:35–8.

22. Goetschmann S, Dibernardo S, Steinmann H, Pavlovic M, Sekarski N, Pfammatter JP, Frequency of severe pulmonary hypertension complicating “isolated” atrial septal defect in infancy. American Journal of Cardiology. 2008; 102: 340–42.

23. Walker RE, Moran AM, Gauvreau K, Colan SD. Evidence of adverse ventricular inter-dependence in patients with atrial septal defects. American Journal of Cardiology. 2004; 93: 1374–77.

24. Masutani S, Senzaki H. Left ventricular function in adult patients with atrial septal defect: implication for development of heart failure after transcatheter closure. Journal of Cardiac Failure. 2011; 17: 957–63.

25. Sachweh JS, Daebritz SH, Hermanns B, et al. Hypertensive pulmonary vascular disease in adults with secundum or sinus venosus atrial septal defect. Ann. Thor.Surg. 2006;81: 207–13.

26. Di Salvo G, Drago M, Pacileo G, et al. Atrial function after surgical and percutaneous closure of atrial septal defect: a strain rate imaging study. Journalof American Society for Echocardiography. 2005; 18: 930–33.

27. Sugimoto M, Ota K, Kajihama A, Nakau K, Manabe H, Kajino H. Volume overload and pressure overload due to left-to-right shunt-induced myocardial injury. Evaluation using a highly sensitive cardiac Troponin-I assay in children with congenital heart disease. Circulation. 2011; 75: 2213–19.

28. Ching YH, Ghosh TK, Cross SJ, et al. Mutation in myosin heavy chain 6 causes atrial septal defect. Nature of Genetics; 2005; 37: 423–28.

29. Chen Y, Han ZQ, Yan WD, et al. A novel mutation in *GATA4* gene associated with dominant inherited familial atrial septal defect. Journal of Thoracic Cardiovascular Surgery. 2010; 140: 684–87.

30. D’Amato E, Giacopelli F, Giannattasio A. et al. Genetic investigation in an Italian child with an unusual association of atrial septal defect, attributable to a new familial *GATA4* gene mutation, and neonatal diabetes due to pancreatic agenesis. J. Diab. Med. 2010; 27:1195–1200.

31. Bakker MK, Kerstjens-Frederikse WS, Buys CH, de Wall HE, et al. First-trimester use of paroxetine and congenital heart defects: a population-based case-control study. Birth Defects Research and Clinical Molecular Teratology. 2010; 88: 94–100.

32. Miller A, Riehle-Colarusso T, Siffel C, Frías JL, Correa A. Maternal age and prevalence of isolated congenital heart defects in an urban area of the United States. American Journal of Medical Genetics. 2011; 155A: 2137–45.

33. Parker SE, Werler MM, Shaw GM, Anderka M, Yazdy MM. National Birth Defects Prevention Study.Dietary glycemic index and the risk of birth defects. American Journal of Epidemiology. 2012; 176: 1110–20.

34. Hala MB, Ghada MS, Mohammed A, Alrefaey A. Atrial septal defects: clinical presentation and recent approach in its diagnosis and treatment. Menoufia Medical Journal. 2013; 27:145–151.

35. Amirah MA, Nada MA, Anna A, Mowafa SH, Ashraf E. The epidemiology of congenital heart diseases in Saudi Arabia: A systematic review. Journal of Public Health Epidemiology; 2015; 7(7): 232–240.

36. Talargia F, Seyoum G, Moges T. Congenital heart defects and associated factors in children with congenital anomalies. Ethiop Med Journal. 2018; vol. 56, no. 4.

37. Benjamin EJ, Muntner P, Alonso A, Bittencourt MS, Callaway CW, Carson AP, et al. Heart disease and stroke statistics - 2019 update: a report from the American Heart Association; on behalf of the American Heart Association Council on Epidemiology and Prevention Statistics Committee and Stroke Statistics Subcommittee. [published online ahead of print January 31, 2019]. Circulation. 2019: 10: 1–3.

38. Marelli AJ, Mackie AS, Ionescu-Ittu R, Rahme E, Pilote L. Congenital heart disease in the general population: changing prevalence and age distribution. Circulation. 2007; 115:163–72.

39. Kazmouz S, Kenny D, Cao Q, Kavinsky C, Hijazi Z, Transcatheter closure of secundum atrial septal defects. Journ.Invas.Cardiol. 2013; 25(5), 257–264.

40. McGill-Lane S, Cusick M, Burke E, Murphy C. Atrial Septal Defect Guideline: What the Nurse Caring for a Patient with Congenital Heart Disease Needs to Know. 2017; 1–13.

41. Reller MD, Strickland MJ, Riehle-Colarusso T, et al. Prevalence of congenital heart defects in metropolitan Atlanta, 1998-2005. Journal of Pediatrics. 2008; 153:807.

42. Wu MH, Chen HC, Lu CW, Wang JK, Huang SC, Huang SK. Prevalence of congenital heart disease at live birth in Taiwan. Journal of Pediatrics. 2010; 156:782.

43. Schwedler G, Lindinger A, Lange PE, et al. Frequency and spectrum of congenital heart defects among live births in Germany: a study of competence Network for Congenital Heart Defects. Clinical Research and Cardiology. 2011; 100:1111.

44. van der Linde D, Konings EE, Slager MA, et al. Birth prevalence of congenital heart disease worldwide: a systematic review and meta-analysis. Journal of American Collective Cardiology. 2011; 58:2241.

45. Lee SA, Moon HS., Lavulo, L., Cho, K.O. and Hyun, C. Isolation, characterization and genetic analysis of canine *GATA4* gene in a family of Doberman Pinschers with an atrial septal defect. Journal of Genetics. 2007; 86: 241–47.

46. Bostan OM, Cil E, Ercan I. The prospective follow-up of the natural course of interatrial communications diagnosed in newborns. European Heart Journal. 2007; 28:16: 2001–5.

47. Haque MS, Hilr K, Mahmud RS. Presentation of atrial septal defect: Symptoms and signs. Journal of Dhaka Medical College. 2011; 20(1): 9–11.

48. Ejim EC, Anisiuba BC, Ike SO, Essien LO, (). Atrial septal defects presenting initially in adulthood: patterns of clinical presentation in Enugu, South-East Nigeria. Journal of Tropical Medicine. 2011; 3:2–3.

49. Fauci S, Braunwald E, Kasper L, Longo L, Hauser L, Jameson JL, Loscalzo J. Harrison’s Principles of Internal Medicine. (^18^thed.). Copyright © The McGraw-Hill Companies. Inc; 2012.

50. Le Gloan L, Legendre A, Iserin L, Ladouceur M. Pathophysiology and natural history of atrial septal defect. Journal of Thoracic Disease. 2017; 10(24):S2854–S2863.

51. Andrews R, Tulloh R. Magee A, Anderson D. Atrial septal defect with failure to thrive in infancy: hidden pulmonary vascular disease? Pediatric Cardiology. 2002; 23:528–30.

52. Lammers A, Hager A, Eicken A, Lange R, Hauser M, Hess J. Need for closure of secundum atrial septal defect in infancy. Journal of Thoracic and Cardiovascular Surgery. 2005; 129: 1353–57.

